# Serological Evidence of Widespread Exposure to H5 Avian Influenza Virus in Arctic Foxes in Svalbard, Norway

**DOI:** 10.64898/2026.09.17.752278

**Authors:** Johanna Hol Fosse, Eva Fuglei, Francesco Bonfante, Line Olsen, Luca Bordes, Rebecca Davidson, Torill Mørk, Ida Kristin Myhrvold, Lone T. Engerdahl, Sandra Venema, Kjersti Sturød, Johan Åkerstedt, Olav Hungnes, Monika Z. Ballmann, Lineke Begeman, Ingebjørg Helena Nymo, Ragnhild Tønnessen, Bjørnar Ytrehus

## Abstract

The continued circulation of H5 clade 2.3.4.4b highly pathogenic avian influenza virus (HPAIV) has caused extensive mortality in wild bird populations worldwide with increasing spillover to mammals. In 2022, H5 HPAIV emerged in Svalbard, Norway, with subsequent detections in wild birds, walruses, polar bears, and arctic foxes. To understand the population-level exposure among Svalbard arctic foxes, we analysed body fluids from carcasses trapped in 2006-2015 (n = 56), 2023-2024 (n = 112), and 2024-2025 (n = 94) for antibodies to H5 avian influenza (anti- H5), influenza A nucleoprotein (anti-NP), and neuraminidase subtypes using ELISAs, haemagglutination inhibition (HI), and a multiplex assay. Only three samples from 2006-2015 tested positive for anti-H5 and were interpreted as false positives. In 2023-2024, seropositivity for anti-H5 was high (95%), supported by a lower anti-NP seropositivity (85%) and antibody profiles consistent with mixed H5N1 (42%) and H5N5 (52%) exposure. In 2024-2025, anti-H5 and anti-NP seroprevalences remained high (83% and 57%), with H5N5 (87%) exposure predominating over H5N1 (3%). A subset of anti-H5-positive samples tested positive by HI (2023-2024: 30%; 2024-2025: 14%). Juveniles with exposure limited to the previous season had higher odds of anti-H5 seropositivity in 2023-2024 than in 2024-2025 (OR 5.5). Analysis of paired lung extracts from a subset of individuals (n = 63) yielded results concordant with body fluids, using an indirect anti-H5 ELISA adapted for carnivores. Our findings demonstrate widespread H5 virus exposure. Together with occasional reports of progression to fatal HPAI, this highlights the need for continued population monitoring to evaluate ecological consequences.

## Introduction

During 2021-2022, Europe experienced an unprecedented epidemic of H5 highly pathogenic avian influenza (HPAI) [1]. Since then, H5 clade 2.3.4.4b HPAI viruses (HPAIV) have become established in wild bird populations, causing recurrent mass mortality events [2–5], and increasing spillover into mammals [6]. Because of their substantial impact on wildlife and livestock health, as well as significant zoonotic potential, understanding H5 HPAIV transmission dynamics and spillover to mammals is essential within a One Health framework [7].

The High Arctic archipelago of Svalbard lies between mainland Norway and the North Pole, at the intersection of multiple migratory flyways. Each spring, large numbers of seabirds, waders, ducks, three goose species, and one passerine (snow bunting, *Plectrophenax nivalis*) migrate to the archipelago to breed before returning to lower latitudes in autumn [8–10]. These seasonal movements create opportunities for repeated introductions of avian influenza viruses from geographically distant regions. The marine ecosystem also supports Atlantic walrus (*Odobenus rosmarus*) and polar bear (*Ursus maritimus*) populations. In contrast, Svalbard’s terrestrial vertebrate community is relatively simple and consists of only a few resident species, including arctic foxes (*Vulpes lagopus*), reindeer (*Rangifer tarandus platyrhynchus*), Svalbard rock ptarmigan (*Lagopus muta hyperborea*), and the introduced Eastern European vole (*Microtus levis*) [11,12].

HPAIV was detected in Svalbard for the first time in 2022 [13]. Both H5N1 and H5N5 viruses were detected among seabirds in 2022-2023 [13], with substantial impact on breeding populations, particularly great skuas (*Stercorarius skua*) [14–16]. During 2023, detections expanded to additional seabird species and Atlantic walrus, while no detections were made in 2024 [13,17]. More recently, HPAIV-associated mortality was documented in a second walrus and a polar bear in Svalbard, while serological investigations revealed widespread exposure among Svalbard polar bears [18].

The arctic fox is the only resident terrestrial predator in Svalbard and occupies a central position in both marine and terrestrial food webs [19–21]. As an opportunistic predator and scavenger feeding on seabirds, seal carrion, tundra birds, and reindeer carcasses, it is likely to encounter HPAIV through multiple pathways. Consequently, arctic foxes may be highly exposed to HPAIV and could serve as useful sentinels for monitoring virus circulation in remote Arctic ecosystems. In 2025, HPAI-associated neurological disease was detected in arctic fox pups from two locations in Svalbard, providing direct evidence that infection can occur and occasionally result in severe disease and mortality [13]. Despite increasing evidence of HPAIV circulation in Arctic wildlife, population-level exposure, survival following infection, and temporal patterns of virus circulation among wild arctic foxes remain unknown.

The aim of this study was to estimate exposure of the Svalbard arctic fox population to H5 avian influenza viruses using serological analyses of carcass-derived samples collected during annual trapping. We analysed samples from arctic foxes trapped before (2006-2015) and after (2023- 2025) the emergence of HPAIV in Svalbard to assess temporal variation in seropositivity and exposure to different neuraminidase subtypes. Our findings provide a pre-HPAIV serological baseline, place the 2025 clinical cases in an epidemiological context, inform carcass-based surveillance, and contribute to understanding HPAIV circulation in Arctic ecosystems.

## Materials and methods

### Arctic fox trapping and necropsy

The arctic fox population in Svalbard is abundant, stable, and classified as being of *Least Concern* [22], with limited licenced trapping by permanent Svalbard residents. All animals included in the study were legally harvested under the Svalbard Environmental Protection Act and Regulation About Harvesting on Svalbard (authors’ translation) [23].

The Norwegian Polar Institute (NPI) monitors the arctic fox population in Svalbard, partly through carcasses collected during annual trapping in Nordenskiöld Land, Spitsbergen. Trapped arctic fox carcasses are frozen at -80° C for five to seven days to eliminate any infective *Echinococcus multilocularis*, thereafter stored at -20° C, skinned, refrozen, and shipped to the Norwegian Veterinary Institute (NVI) in Tromsø for necropsy and sample collection.

Carcasses were weighed, sexed, and assigned a body condition score using a subjective fat index: 0 (no visible fat), 1 (barely measurable fat), 2 (subcutaneous fat deposits over the rump and intraabdominal fat), 3 (subcutaneous fat deposits over the rump, belly, and flanks and intraabdominal fat), or 4 = (subcutaneous fat deposits covering most of the body and intraabdominal fat) [24]. Age was estimated by counting cementum annuli in sectioned canine teeth, with the term “juvenile” used to describe individuals born the previous spring [25]. Metadata for all individuals are provided in Table S1.

### Nucleic acid extraction and molecular screening

Tissue samples from individuals trapped in 2024-2025 were screened for rabies virus (brain, n = 97) and influenza A virus (brain and tracheal swabs, n = 97). Nucleic acids were extracted using the automated MagNA Pure 96 system (Roche) in accordance with the manufacturer’s instructions. Detection of influenza A virus was performed using a real-time reverse transcription–PCR (RT-PCR) assay targeting the matrix (M1) gene, as described [26]. For rabies virus detection, a real-time RT-PCR assay targeting a conserved region of the nucleoprotein (N) gene was used, following established protocols [27]. One individual was excluded from the study due to detection of rabies virus, while two carcasses did not yield body fluids suitable for analysis, resulting in 94 carcasses progressing to serological analyses.

### Serological sample collection and processing

Available body fluid samples from individuals trapped or found dead between 2006 and 2015 (n = 56) were included to establish a baseline seroprevalence and evaluate assay specificity. Samples for serological analysis were collected from foxes trapped in 2023-2024 (n = 112) and 2024-2025 (n = 94).

Body fluid was collected from the thoracic or abdominal cavity, clarified by centrifugation (2300 ×*g*, 15 min, room temperature), and stored at -20 °C. After thawing, fluids were heat-inactivated (56 °C, 30 min) and re-centrifuged (11,000 ×*g*, 10 min, 4 °C). Lung extracts, increasingly used in serological surveillance of wild carnivores as a practical alternative to the collection of thoracic fluids, were prepared from paired lung samples collected from 63 individuals trapped in 2023-2024 and stored at -20 °C. After thawing, a tissue piece (∼ 0.5 cm x 0.5 cm x 2.5 cm) was excised, transferred to an Eppendorf tube containing 1 mL phosphate-buffered saline, and incubated at room temperature for 22 min, including 4 min of agitation (500 rpm). The tissue was then removed, and the supernatant was heat-inactivated (56 °C, 30 min) and centrifuged (500 ×*g*, 10 min, 4 °C).

### Enzyme-linked immunoassays (ELISAs)

Antibodies to H5 avian influenza (anti-H5) were detected by the ID Screen Influenza H5 Antibody Competition 3.0 Multi-species ELISA kit (Innovative Diagnostics; anti-H5 cELISA), used according to the manufacturer’s instructions for fox serum (Addendum FLUACH5V3 ver 150725 EN). Samples with signal/noise (S/N)-ratio ≤ 40% were defined as positive. Complementary detection of anti-H5 was performed by the ID Screen Influenza H5 Indirect ELISA kit (Innovative Diagnostics; anti-H5 iELISA), used according to the manufacturer’s instructions (FLU5S ver 0123 EN) with modifications to allow use in arctic foxes. Briefly, samples were diluted 1:100 and incubated (60 min, 21 °C). After washing, an anti-dog horseradish peroxidase-conjugated antibody diluted 1:10 in buffer 1 (both: Innovative Diagnostics) was added and incubated (30 min, 21 °C). Substrate and stop solutions were added according to the manual. Based on prior assay optimization carried out at the Istituto Zooprofilattico Sperimentale delle Venezie (IZSVe, data not shown), samples with optical density (OD) > 0.09 were defined as positive.

Antibodies to influenza A nucleoprotein (anti-NP) were detected by the ID screen Influenza A Antibody Competitive Multi-species (Innovative Diagnostics; anti-NP cELISA) was performed according to the manufacturer’s instructions for ferret, cat, and dog serum (Addendum FLUACA ver 0917 EN (04/2025)). Samples with S/N-ratio ≤ 45% were defined as positive.

### Haemagglutination inhibition tests

Haemagglutination inhibition (HI) tests were performed in accordance with the WOAH Terrestrial manual [28] and the guidelines for diagnosis of H5Nx HPAI virus infection in mammals provided by IZSVe [29]. Samples with HI titres ≥20 were defined as positive. Despite pre-treatment with chicken erythrocytes, a subset of samples caused autoagglutination and were excluded from the analysis. Anti-H5-positive samples from 2006-2015 and 2023-2024 were tested with two antigens: one representing currently circulating H5 clade 2.3.4.4b viruses (A/turkey/Italy/VIR9520-320/2021(H5N1)), and one representing low pathogenic H5 viruses (A/teal/England/7394/2006 (H5N3)). Samples from 2024-2025 were only tested with the H5 clade 2.3.4.4b antigen.

### Multiplex serological assay

Antibody binding to haemagglutinin (HA) 1-16 and neuraminidase (NA) 1-9 was analysed using Luminex xMAP technology [30]. Samples were tested against 71 AIV proteins distributed across six multiplex assays. Multiplexes and samples were tested at 1:50 dilution in sample buffer (PBS, 0.05% Tween 20, 10% PRIblocker). Sample specific cut-offs were calculated separately for HA and NA. For HA, the five beads with the highest mean fluorescence intensity (MFI) values were excluded and the cut-off was defined as the mean MFI of the remaining beads plus five standard deviations. The same approach was used for NA, except that the three beads with the highest MFI values were excluded. Positive signals to multiple subtypes were evaluated for potential cross-reactivity. Signals to genetically distinct subtypes were considered evidence of independent antibody responses, as the arctic foxes may have been exposed to multiple influenza A virus subtypes throughout their lifetime. Samples were classified as seronegative when no bead set exceeded both the assay cut-off and the minimum signal threshold of 50 MFI.

### Data processing and statistics

Data from standard diagnostic tests (anti-H5 cELISA, anti-NP cELISA, and HI) were extracted from the NVI laboratory information management system and combined with metadata and results from anti-H5 iELISA and multiplex assay in Microsoft Excel. The complete dataset is provided in Table S1. GraphPad Prism (v10.3.1) was used for graph generation and statistical analysis (Fisher’s exact test and McNemar’s test based on discordant pairs), with statistical significance set at p<0.05. Test specificity and inter-assay agreement were calculated using MedCalc online calculators (https://www.medcalc.org/en/calc/).

## Results

### Molecular screening

One individual tested positive for rabies virus and was excluded from further analyses. No individuals tested positive for influenza A virus.

### Serological screening of body fluids from arctic foxes trapped in Svalbard reveals widespread exposure to H5 avian influenza

Using a commercially available competitive ELISA (anti-H5 cELISA), we detected anti-H5 in a high proportion of samples from arctic foxes trapped after the emergence of H5 HPAIV in Svalbard, with 106 of 112 (95%) individuals from 2023-2024 and 78 of 94 (83%) individuals from 2024-2025 testing positive (Fig. 1A). In contrast, only 3 of 63 (5%) individuals from the pre-emergence period (2006-2015) tested positive. Samples from 2023-2024 and 2024-2025 were also tested for antibodies to influenza A nucleoprotein (anti-NP), revealing positive results in 95 of 112 (85%) and 54 of 94 (57%) individuals, respectively (Fig. 1B). Anti-H5-positive samples were subsequently tested by HI using a H5 clade 2.3.4.4b antigen. HI titres ≥20 were detected in 21 of 69 (30%) samples from 2023-2024 and 10 of 71 (14%) samples from 2024- 2025, but in none of the three samples from 2006-2015 (Fig. 1C). Complementary HI testing of anti-H5-positive samples from 2006-2015 and 2023-2024 using a Eurasian-lineage low pathogenic avian influenza H5 antigen yielded no positive results.

**Figure 1:**
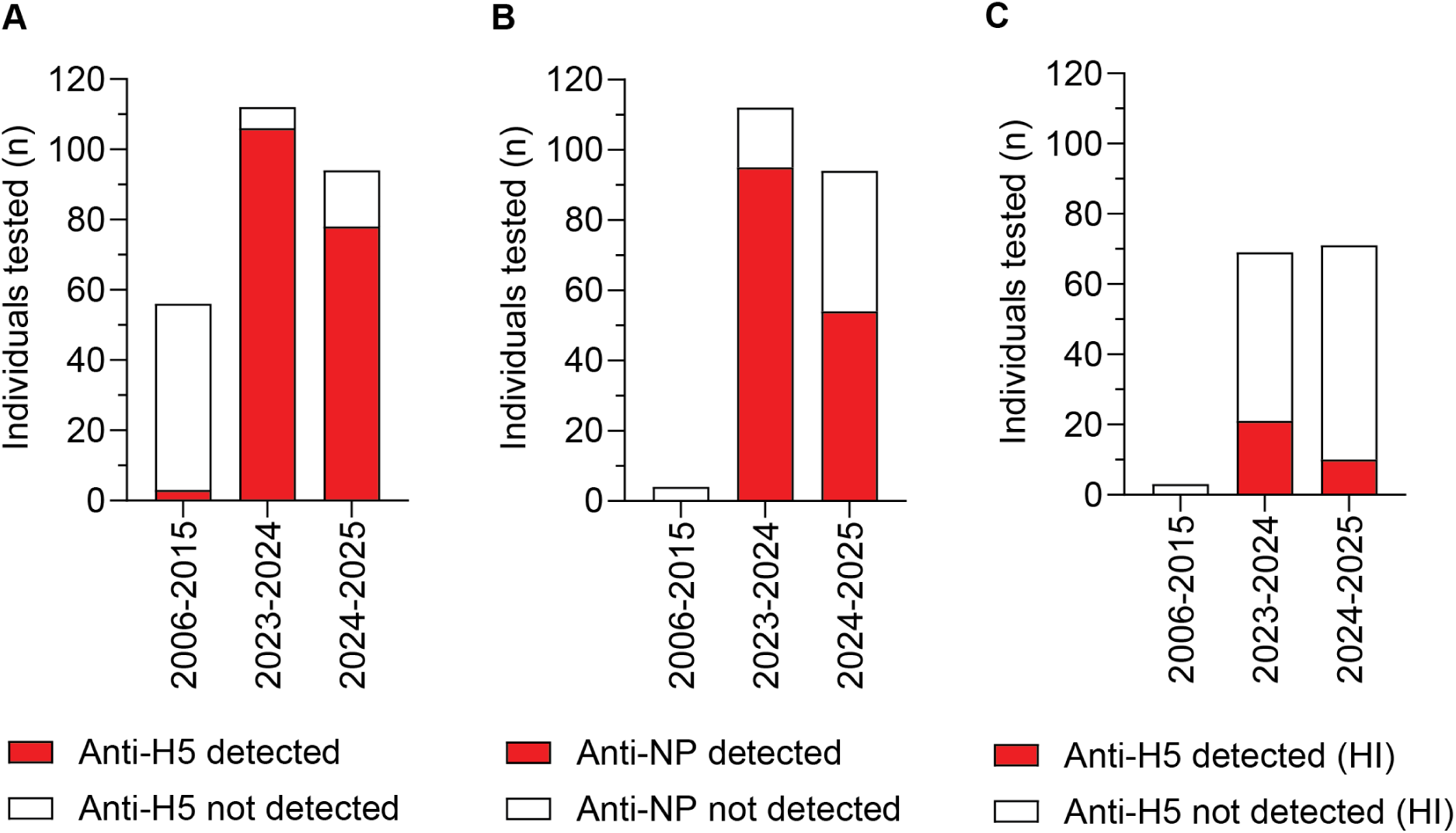
Serological evidence of widespread exposure to H5 avian influenza in Svalbard arctic foxes. Body fluids from arctic fox carcasses obtained in 2006-2015 (n = 56), 2023-2024 (n = 112), and 2024-2025 (n = 94) were analysed for (A) antibodies to H5 avian influenza (anti- H5) and (B) influenza A nucleoprotein (anti-NP) using commercial competitive ELISA kits. (C) Samples tested positive for anti-H5 by ELISA were also tested for haemagglutination inhibition (HI) using a A/turkey/Italy/VIR9520-320/2021(H5N1) antigen.

### Diagnostic performance of different matrix-assay combinations to detect anti-H5 in arctic fox carcasses

Body fluid samples from 2006-2015 (n = 56) and those samples from 2023-2024 with sufficient material (n = 88) were re-analysed using a customized indirect anti-H5 ELISA (anti- H5 iELISA). One sample from 2006-2015 (2%) and 85 samples from 2023-2024 (97%) tested positive (Fig. 2A), showing excellent agreement with the anti-H5 cELISA (95% agreement, Cohen’s kappa = 0.900, Table 1). The four samples from 2006-2015 that tested positive by either anti-H5 cELISA (n = 3) or anti-H5 iELISA (n = 1) all tested negative in the anti-NP cELISA, and three also tested negative by HI (Fig. 1B-C). The fourth contained insufficient material for HI testing. Given the epidemiological context and the lack of concordance among assays, anti-H5-positive results from 2006-2015 were interpreted as false positives. Accordingly, estimated test specificities were 95% (95% CI: 85%-99%) for the anti-H5 cELISA and 98% (95% CI: 90%-100%) for the anti-H5 iELISA, with no significant difference between the assays.

**Figure 2:**
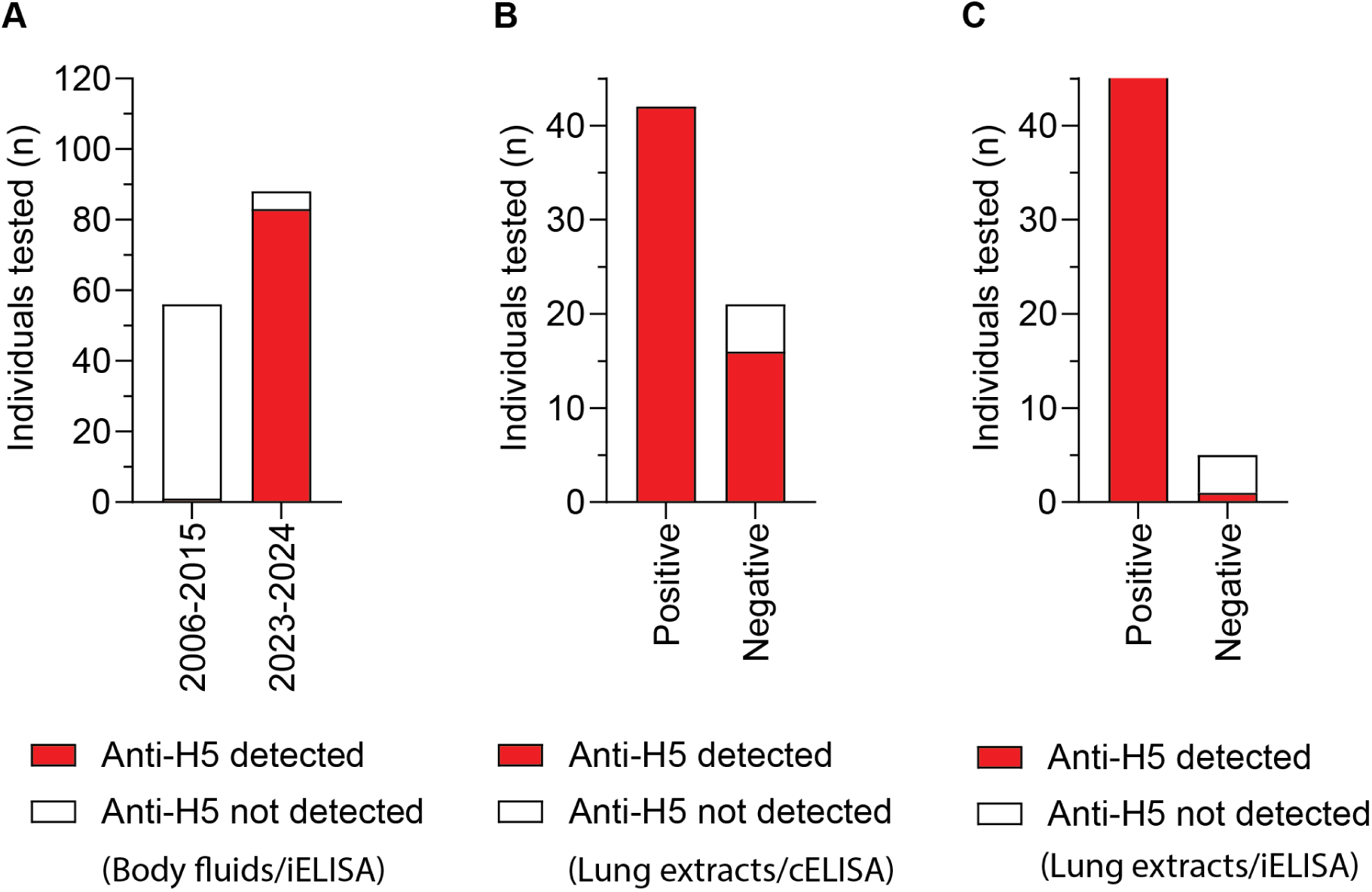
Diagnostic performance of anti-H5 ELISAs in samples from arctic fox carcasses. (A) Body fluids from arctic fox carcasses obtained in 2006-2015 (n = 56) and 2023-2024 (n = 88) were analysed by a customised indirect anti-H5 iELISA. (B-C) Paired lung extracts of arctic fox carcasses from 2023-2024 (n = 63) were analysed by (B) anti-H5 cELISA or (C) anti-H5 iELISA, and results were compared to those obtained from analysing body fluids by anti-H5 cELISA (Positive/Negative). Test agreements are provided in Table 1.

**Table 1.**
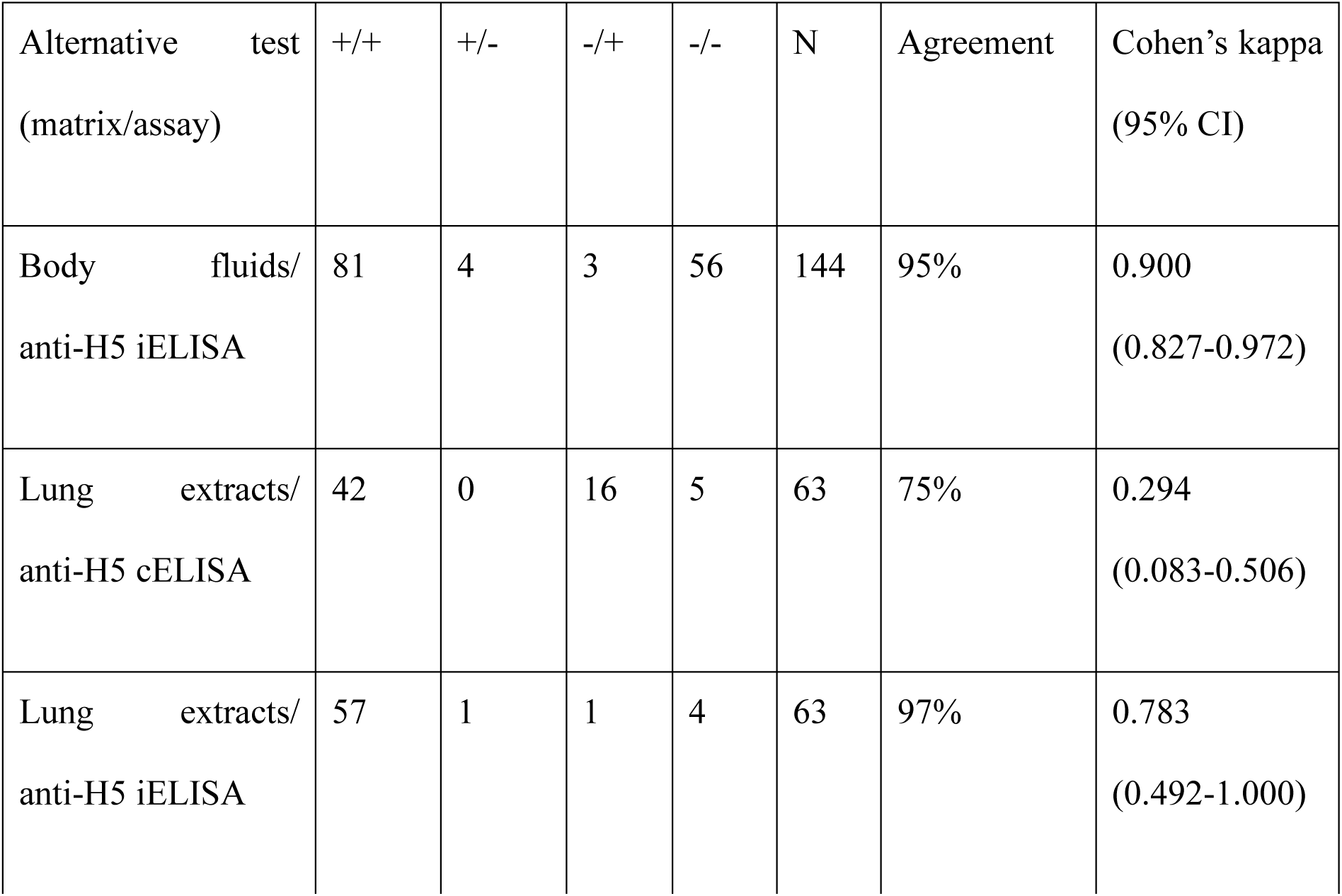
Performance of different matrix-assay combinations for detecting anti-H5 antibodies in samples from arctic fox carcasses. For the result categories (+/+, +/−, −/+, and −/−), the first symbol refers to the anti-H5 cELISA performed on body fluids and the second symbol refers to the comparator test shown in the left column. N denotes the total number of samples tested.

| Alternative test<br>(matrix/assay) | +/+ | +/- | -/+ | -/- | N | Agreement | Cohen's kappa<br>(95% CI) |
| --- | --- | --- | --- | --- | --- | --- | --- |
| Body fluids/<br>anti-H5 iELISA | 81 | 4 | 3 | 56 | 144 | 95% | 0.900<br>(0.827-0.972) |
| Lung extracts/<br>anti-H5 cELISA | 42 | 0 | 16 | 5 | 63 | 75% | 0.294<br>(0.083-0.506) |
| Lung extracts/<br>anti-H5 iELISA | 57 | 1 | 1 | 4 | 63 | 97% | 0.783<br>(0.492-1.000) |

Paired lung extracts from 2023-2024 (n = 63) were tested using both anti-H5 ELISAs (Fig. 2B-C). Apparent seropositivity was higher with the anti-H5 iELISA than with the anti-H5 cELISA (92% versus 67%, exact McNemar’s test, p < 0.005). All 16 discordant samples were positive by the anti-H5 iELISA and negative by the anti-H5 cELISA, indicating systematic directional disagreement between the assays. Results from analysing lung extracts by anti-H5 iELISA showed good agreement with results from paired body fluid samples analysed by anti- H5 cELISA (97% agreement, Cohen’s kappa = 0.783, Table 1).

### Geographical distribution of trapping sites

No geographical clustering of anti-H5-positive individuals was observed (Fig. 3).

**Figure 3:**
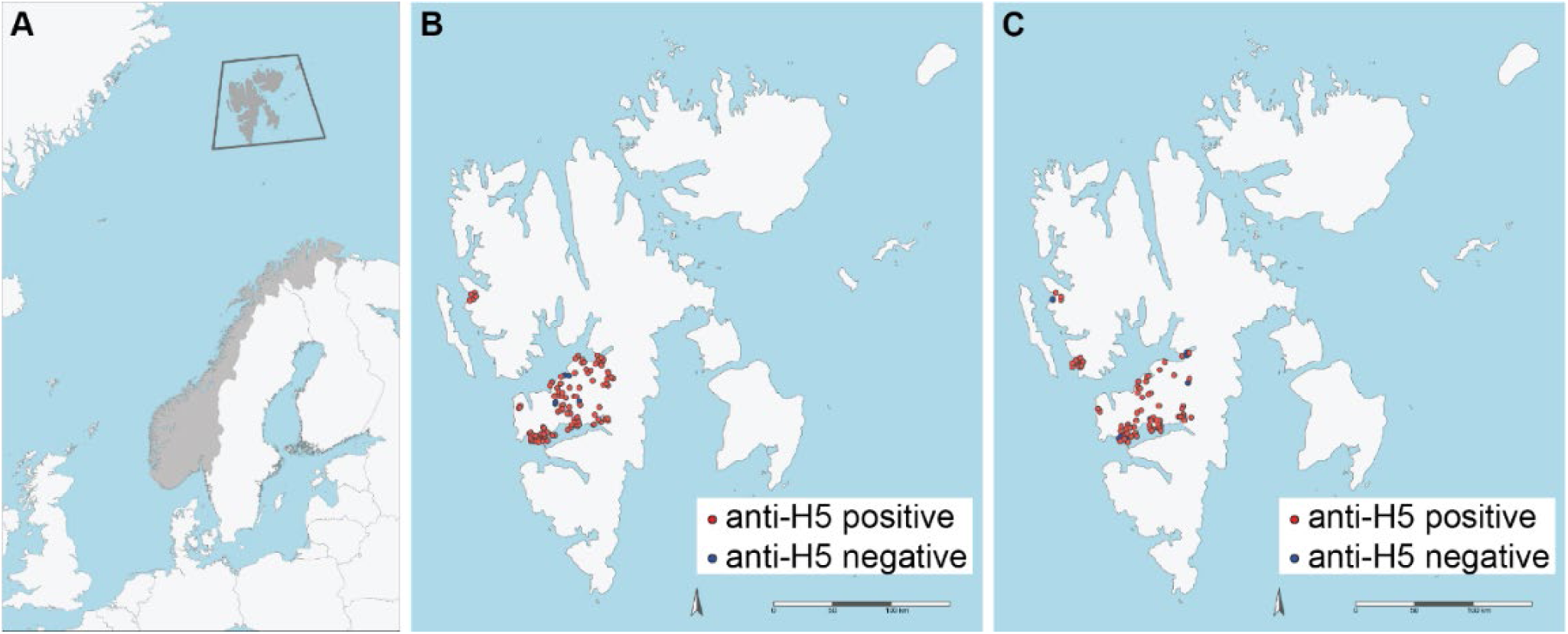
Geographical distribution of arctic fox trapping sites in Svalbard in 2023-2024 and 2024-2025. (A) Map of Norway (grey). The boxed area marks the geographical location of the Svalbard archipelago. (B-C) Location of trapping sites at the Svalbard archipelago in 2023-2024 (B) and 2024-2025 (C). Scale bars represent 150 km.

### Age-guided comparison of serological evidence of exposure in 2023-2024 and 2024-2025

Assuming exposure risk is highest during summer, when large numbers of seabirds breed in Svalbard, we focused on juveniles born the previous spring, whose exposure would largely be limited to the preceding season. Most trapped individuals were young, with 81 juveniles identified in 2023-2024 (Fig. 4A) and 63 in 2024-2025 (Fig. 4B). Juveniles trapped in 2023- 2024 had approximately five-fold higher odds of testing positive by anti-H5 cELISA than those trapped in 2024-2025 (OR 5.50, 95% CI: 1.72-15.88, Fisher’s exact test, p<0.01, Fig. 4C). No significant differences were identified between juveniles and older (2–9-year-old) individuals (Fig. 4D) or between females and males across seasons (Fig. 4E). However, emaciated individuals (fat index = 1) had lower odds of testing positive by anti-H5 cELISA (OR 0.29, 95% CI 0.12-0.76, Fisher’s exact test, p<0.05, Fig. 4F).

**Figure 4:**
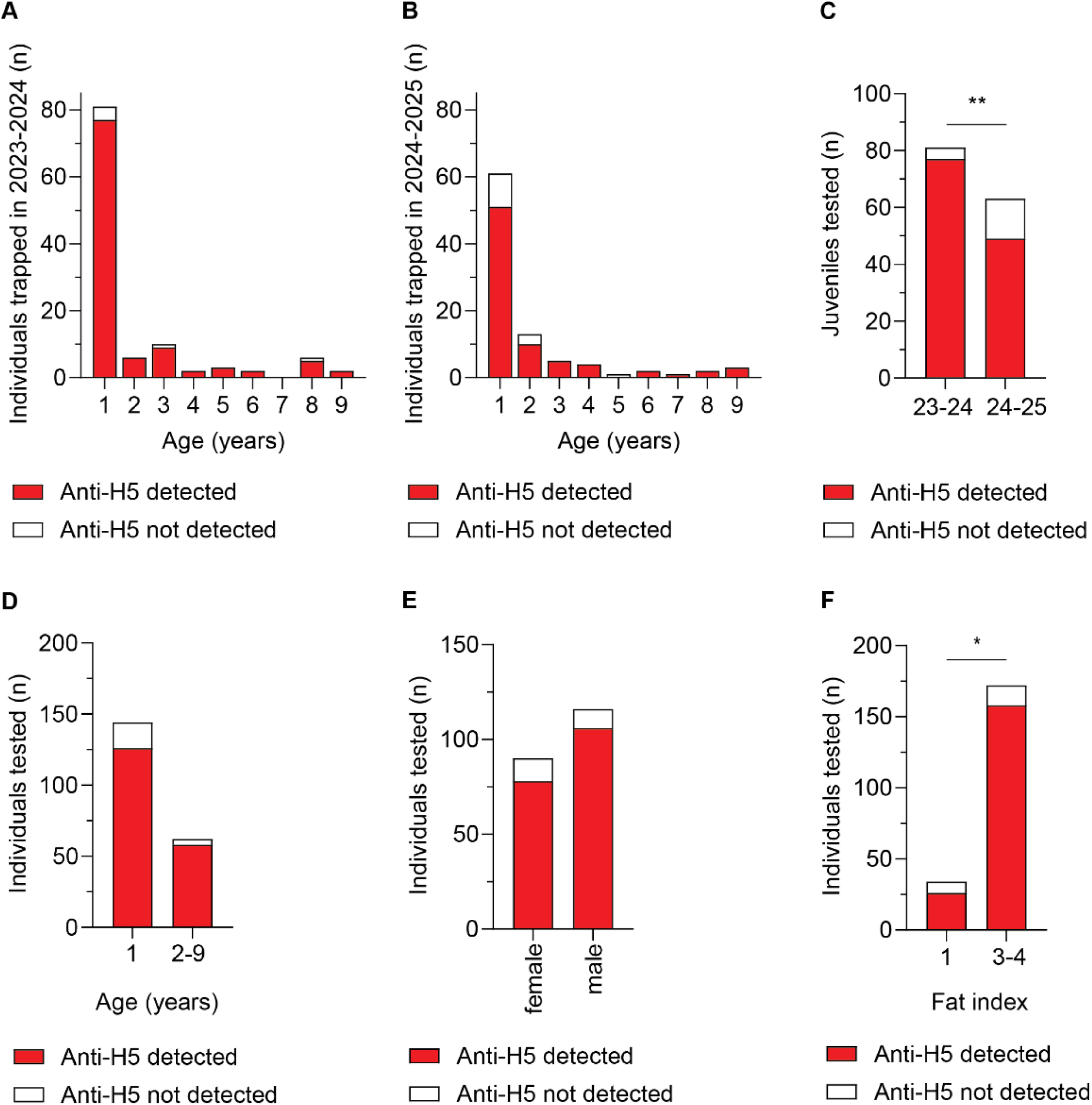
Age-guided comparison of seroprevalence to H5 avian influenza in Svalbard arctic foxes trapped in 2023-2024 and 2024-2025. (A-D) Cementum annuli counts were used to estimate the age of trapped individuals from (A) 2023-2024 and (B) 2024-2025 and compare the detection of anti-H5 antibodies in body fluids by a commercial competitive ELISA. (C) Juveniles from 2023-2024 were more likely to be seropositive than juveniles from 2024-2025 (**p<0.01, Fisher’s exact test, OR 5.50). (D) Juveniles did not show different odds for testing positive than 2-9-year-old individuals, across years. (E) Females and males did not show different odds for testing positive across years. (F) Emaciated individuals (fat index = 1) showed lower odds of testing positive than individuals with fat index 2-4 (*p<0.05, Fisher’s exact test, OR 0.29).

### Antibody profiles indicate a shift from mixed avian influenza subtype exposure in 2023-2024 to predominantly H5N5 exposure in 2024-2025

To investigate which H5 avian influenza strains contributed to exposure in Svalbard arctic foxes, 60 anti-H5-positive samples with low S/N-values from each trapping season were analysed using a multiplex assay for antibodies to avian influenza subtypes [30]. Anti-HA was detected in 45 samples from 2023-2024 and 46 samples from 2024-2025, consistent with the limited sensitivity of the assay. Anti-H5 predominated in all positive samples (Table S1). Among samples from 2023-2024, anti-N1 and anti-N5 were detected at similar frequencies (Fig. 5A). Anti-N1 predominated in 26 samples (43%), 12 showing concurrent reactivity to N5. Anti-N5 predominated in 31 samples (52%), including 12 with concurrent N1 reactivity, one of which also reacted with N7. Three samples lacked detectable anti-NA. This pattern shifted in 2024-2025, when samples predominantly displayed anti-N5 (Fig. 5A, Table S1). Anti-N1 dominated in only 3 samples (5%), one of which also displayed reactivity to N5, while anti-N5 dominated in 51 samples (85%), of which eight also reacted with other subtypes (N1, n = 7, N8, n = 1). Six samples tested negative for anti-NA. The three individuals from 2024-2025 that predominantly displayed anti-N1 were estimated to be 3, 3, and 4 years old (Table S1). In contrast, all one-year-olds from 2024-2025 predominantly displayed anti-N5 reactivity (Fig. 5B, Table S1). The odds of predominant anti-N5 reactivity were significantly higher in 2024- 2025 than 2023-2024, both overall and when the analysis was restricted to one-year olds (Fisher’s exact test, p<0.0001).

**Figure 5:**
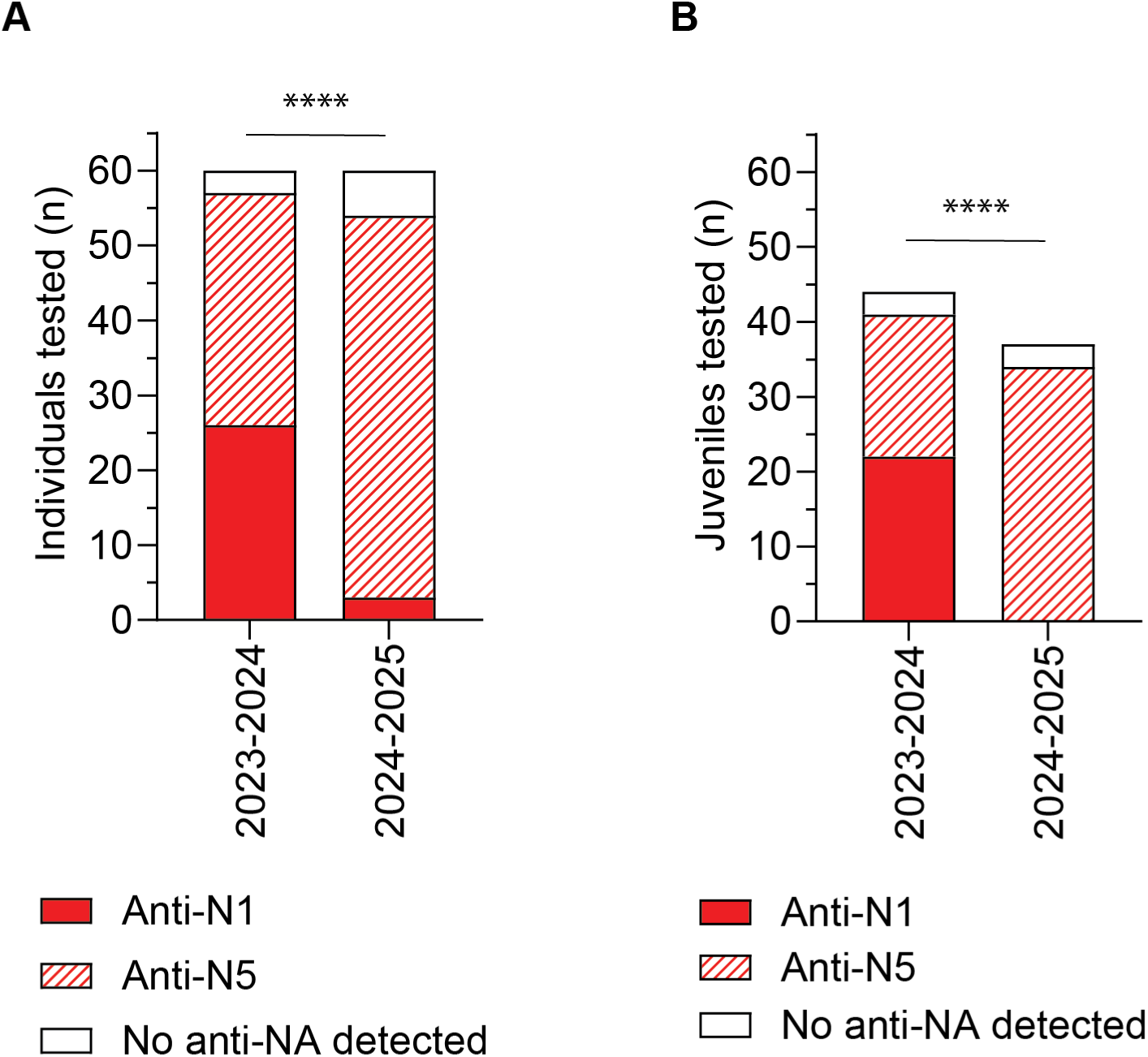
Antibody profiles reveal a shift from mixed avian influenza subtype exposure in Svalbard arctic foxes trapped in 2023-2024 to predominantly H5N5 exposure in 2024-2025. Anti-H5-positive body fluid samples collected from arctic fox carcasses trapped during 2023-2024 (n = 60) and 2024-2025 (n = 60) were analysed using a multiplex assay designed to distinguish antibodies against influenza A virus neuraminidase (NA) subtypes. (A-B) Bar plots show neuraminidase subtype reactivity profiles for (A) all samples and (B) juvenile foxes only (n = 81). Samples were classified according to predominant antibody reactivity to N1 (anti-N1), N5 (anti-N5), or the absence of detectable anti-NA antibodies. A subset of samples exhibited additional reactivity to other NA subtypes, as described in the main text.

## Discussion

Our study reveals widespread exposure to H5 avian influenza in Svalbard arctic foxes trapped during the winters 2023-2024 and 2024-2025, with no evidence of H5 exposure during the pre-emergence period (2006-2015), as the few anti-H5-positive results detected in that period were interpreted as likely false positives. Antibody profiles changed over time, suggesting a transition from mixed H5N1 and H5N5 exposure in 2023-2024 to predominantly H5N5 exposure in 2024-2025. Our findings demonstrate that many arctic foxes survive exposure to H5 avian influenza viruses and develop a detectable humoral response. However, recent detections of fatal HPAI in arctic fox pups suggest that the virus may have profound effects in some individuals, although its potential effects on population dynamics remain unknown.

The temporal changes in antibody profiles should be viewed in the context of HPAIV detections in Svalbard wildlife. Both H5N1 and H5N5 viruses were detected in birds and marine mammals in 2023, whereas no detections were reported in 2024. However, H5N5 was subsequently detected in arctic foxes in 2025 and in a polar bear and an Atlantic walrus in 2026 [13,18]. Together with widespread H5 seropositivity in polar bears, these findings suggest sustained circulation of H5 viruses within the Svalbard ecosystem and repeated spillover into mammalian hosts, despite relatively few reported detections.

The anti-H5 seropositivity observed in our study is remarkably high. We detected anti-H5 in 95% of foxes trapped in 2023-2024 and 83% of foxes trapped in 2024-2025. The seroprevalence of anti-NP antibodies was slightly lower (85% and 57%). These percentages are much higher than those previously reported in the closely related red fox (*Vulpes vulpes*), where H5 seroprevalences ranged from 16% to 37% in studies from the Netherlands, Ireland, and Germany [31–33]. In Germany, seropositivity was associated with proximity to waterbird-rich habitats, supporting exposure through contact with infected wild birds or their carcasses. In contrast, no seropositive foxes were identified in a survey from Pennsylvania, USA [34]. Together, these findings indicate substantial geographic variation in H5 virus exposure among wild fox populations and suggest exceptionally intense exposure pressure in Svalbard following the emergence of H5 HPAIV.

Given the opportunistic predatory and scavenging behaviour of the arctic fox, together with numerous mortality events and H5 HPAIV detections reported in Svalbard seabirds and marine mammals after 2022, ingestion of infected prey or carrion appears to be the most plausible source of H5 avian influenza exposure [14–17,20,35], although transmission between foxes cannot be excluded. Scavenging by foxes may contribute to ecological resilience by facilitating the removal of HPAI-infected carcasses from the environment [36]. Furthermore, experimental studies suggest that infection acquired through scavenging may result in milder disease than intratracheal infection [37]. Nevertheless, recent documentation of fatal infections in arctic fox pups from two different locations in 2025 [13] raises the possibility that pups could be more vulnerable to severe disease than adults, highlighting the need to assess potential impacts on juvenile survival and recruitment. While denning survival is high in Svalbard, survival through the first winter is low (26%), compared with annual adult survival of 68% [22,38]. HPAIV may therefore represent an additional pressure on the population alongside other threats, including infestation with the blood-sucking louse (*Linognathus* sp.) and occasional rabies outbreaks [39,40]. However, its effects on survival and population dynamics remain unknown.

Seabirds following different migratory flyways and breeding at Svalbard during the summer season [8–10] may serve as a convergence point for avian influenza strains from geographically distinct regions, potentially facilitating viral reassortment [13]. Avian influenza surveillance in Svalbard is currently passive, based on analysis of samples from sick or dead birds and mammals that are submitted for analysis. Limited submission from Svalbard likely underestimates HPAIV circulation in the high Arctic, as illustrated by the discrepancy between reported detections and the high seroprevalence observed in both arctic foxes and polar bears. Our study therefore suggests that serological analyses of trapped arctic foxes may offer useful complementary information about circulating viruses, particularly when focusing on one-year-old individuals with a temporally limited exposure. The antibody profiles observed in our study shifted from 2023-2024 to 2024-2025. Together with the age-stratified analysis, they indicate a slight reduction in total exposure to H5 avian influenza and a clear shift towards the H5N5 subtype. These results demonstrate the potential value of including serological analysis of apex predators to H5 HPAIV surveillance programs in remote locations such as Svalbard. Our study also informs the choice of assay when designing such wildlife serological surveillance programs, by describing the performance of two anti-H5 ELISAs in the analysis of samples from arctic fox carcasses, both of which can be readily implemented by any laboratory equipped for standard serological analyses.

Our study is not designed to determine whether transmission has occurred within the Svalbard arctic fox population, which would require detection of a higher number of viral genomes than what has been reported [13]. The lack of influenza A virus RNA in tracheal swabs and brains of carcasses from 2024-2025 is consistent with a relatively low likelihood of active virus shedding from arctic foxes during winter trapping. This is supported by experimental data from red foxes suggesting that active virus shedding is limited to approximately five to seven days [37]. Exposure pressures are therefore likely to be highest in the summer, when potentially infected seabirds and their carcasses are most accessible as a dietary source. This is temporally separated from the period when most human contact occurs, i.e. during trapping in November to March. Nonetheless, the potential for zoonotic infection should not be ignored, particularly as virus could persist in frozen carcasses. Strict biosecurity measures for those handling arctic fox carcasses are already recommended due to the risk of rabies infection, but future risk assessment revisions may also need to consider the possibility of respiratory exposure.

In conclusion, our findings demonstrate widespread exposure of Svalbard arctic foxes to H5 avian influenza virus following the emergence of clade 2.3.4.4b viruses in the High Arctic, with unknown consequences for the population. The exceptionally high seroprevalence, combined with temporal changes in antibody profiles, indicates substantial and evolving exposure within the Svalbard ecosystem. Arctic foxes may represent a useful sentinel species for monitoring HPAIV circulation in remote regions where conventional surveillance is challenging. Continued integrated surveillance of wildlife hosts, predators, and scavengers will be important for understanding the long-term ecological consequences of HPAIV circulation and for detecting future changes in virus distribution, subtype composition, and transmission dynamics [18].

## Data availability

The raw data that supports the findings of this study are available in Table S1.

## Acknowledgements

The authors would like to thank Emma Rakvåg Vangen, Anna Galina Henriksson, Kristin Ruså Sørby, Frieda Betty Ploss, and Irene Haugen (NVI) for excellent technical assistance. We also thank the local trappers in Svalbard for providing the arctic fox carcasses for research and monitoring, the Governor of Svalbard, and the NPI’s logistic department in Longyearbyen, Svalbard, for the work with collection and storing of the fox carcasses after the trapping.

## Author contribution

Conceptualisation: JHF, LO, RD, RT, BY; Methodology: JHF, EF, FB, LB, IKM, LTE, MZB; Validation: JHF, FB, LB; Formal analysis: JHF, FB, LB; Investigation: JHF, EF, LO, RD, FB, LB, SV, TM, IKM, LTE, KS; Resources: EF, FB, LB, TM, BY; Data curation: JHF, EF, FB, LB; Writing – original draft: JHF; Writing – Review & Editing: All authors; Visualisation: JHF, JÅ; Project administration: EF, FB, LB, RT, BY; Funding acquisition: EF, FB, LB, MZB, RT, BY

## AI declaration

Microsoft Copilot (Microsoft 365 Copilot) was used to improve language, clarity, and readability. All scientific content, interpretation, and conclusions are the responsibility of the authors.

## Appendices

Table S1. Merged test results and metadata from all tested individuals.

